# Nano-scale viscometry reveals an inherent mucus defect in cystic fibrosis

**DOI:** 10.1101/2024.03.06.583796

**Authors:** Olga Ponomarchuk, Francis Boudreault, Ignacy Gryczynski, Sergei V. Dzyuba, Rafal Fudala, Zygmunt Gryczynski, John W. Hanrahan, Ryszard Grygorczyk

**Affiliations:** Centre de recherche du Centre hospitalier de l’Université de Montréal (CRCHUM), Montréal, Québec, Canada; Texas Christian University, Department of Physics and Astronomy, Fort Worth, TX, 76129, USA; Texas Christian University, Department of Chemistry and Biochemistry, Fort Worth, TX, 76129, USA; University of North Texas Health Science Center, Department of Microbiology, Immunology and Genetics, Fort Worth, TX, 76107, USA; Department of Physiology, McGill University, Montreal, Quebec, Canada; Cystic Fibrosis Translational Research Centre, McGill University, Montreal, Quebec, Canada; Département de Médecine, Université de Montréal, Montréal, Québec, Canada

**Keywords:** mucus, mucin granule, cystic fibrosis, viscosity, molecular viscometer

## Abstract

Abnormally viscous and thick mucus is a hallmark of cystic fibrosis (CF). How the genetic defect causes abnormal mucus in CF remains unanswered and a question of paramount interest. Mucus is produced by hydration of gel-forming mucin macromolecules that are stored in secretory granules prior to release. Current understanding of mucin/mucus structure before and after secretion remains limited and contradictory models exist. Here we used a molecular viscometer and fluorescence lifetime imaging of primary epithelial cells (Normal and CF) to measure nanometer-scale viscosity. We found significantly elevated intraluminal nanoviscosity in a population of CF mucin granules, indicating an intrinsic, pre-secretory, mucin defect. Validation experiments showed that high nanoviscosity in cellular environments is mainly due to the low mobility of water that hydrates macromolecules. Nanoviscosity influences protein conformational dynamics and function. Its elevation along the protein secretory pathway indicates molecular overcrowding and is expected to alter mucin’s post-translational processing, hydration, and mucus rheology after release. The nanoviscosity of extracellular CF mucus was elevated compared to non-CF mucus. Remarkably, it was higher after secretion than in granules, which suggests mucins have a weakly-ordered state in granules and adopt a highly-ordered, nematic crystalline structure extracellularly. This challenges the classical view of mucus as a porous agarose-like gel and suggests an alternative model for mucin organization before and after secretion. Our study also suggests that endoplasmic reticulum stress due to molecular overcrowding contributes to mucus pathogenesis in CF cells. It encourages the development of therapeutics that target pre-secretory mechanisms in CF and other muco-obstructive lung diseases.

---

Gel-like viscoelastic mucus plays an essential role in protecting the airways from environmental pathogens, pollutants, dehydration, and shear stress ^1^. Mucus is produced by airway surface mucous/goblet cells and submucosal glands and forms a discontinuous layer covering epithelial surfaces. It entraps inhaled particles and pathogens and helps to remove them from the airways by mucociliary clearance (MCC) mechanism, which is driven by coordinated beating of motile cilia ^2^. MCC critically depends on the composition and physical properties of mucus. Healthy airway mucus contains ∼98% water, polymeric mucins, salts and other constituents (DNA, lipids, cellular debris). On the microscopic scale mucus is often considered a gel resembling agarose, with microscopic domains (i.e., pores) between entangled polymeric mucin fibers that are filled with a low viscosity fluid. The gel-forming polymeric mucins MUC5B, MUC5AC are the main structural components of mammalian respiratory mucus and the largest glycoproteins in the body (5-50 MDa). After translation at the endoplasmic reticulum (ER), mucin macromolecules undergo multistep polymerization/assembly and extensive posttranslational processing in the ER and Golgi. Due to their enormous size this process induces a stress response in the ER ^3^. Glycans constitute up to 80% of the molecular mass of mature, fully glycosylated mucins. Many of the terminal sugars have sulfation/sialylation that cause mucins to have a net negative surface charge. Repulsion between these charges increases the rigidity of the polymer and correlates with mucus viscosity and elasticity ^4^.

Before secretion, partially hydrated mucins are densely packed within intracellular storage granules ^5^, however the intragranular organization of mucins is not fully understood. It is often assumed that they are packed in a well-ordered state that allows a high degree of condensation, that is also favored by charge shielding and ionic crosslinking by high intragranular concentrations of Ca^2+^ and H^+^ ^6^. Well-structured packing may also minimize knots and entanglements of mucin chains that might impair rapid release and expansion ^7, 8^. Hypotheses for mucin organization within granules propose concatenated rings for intestinal mucin MUC2, and crystalline arrays of folded block copolymers for airway MUC5B/5AC. These models envision net-like gel and nematic crystalline-ordered structures, respectively, and predict different swelling dynamics after release ^9, 10^. It has been suggested that two phases coexist within the mucin granule, a mobile fraction and a condensed matrix consisting of mucins that are immobilized by pH-dependent, noncovalent interactions ^11^. Stored mucins are released from airway epithelial cells by exocytosis of mucin granules ^12^. During release the mucins undergo rapid unfolding (<1-s time scale) and expansion in the extracellular space. In healthy airways, this release is accompanied by significant hydration and the mucins expand up to 100-fold through mechanisms that depend on Ca^2+^ chelation and alkalinization by bicarbonate in the extracellular milieu ^13^.

CF lung disease is characterized by abnormally viscous, poorly transportable mucus. It accumulates due to impaired mucociliary clearance, leading to respiratory tract infection, chronic inflammation and airway obstruction. These abnormalities have been attributed to reductions in airway surface liquid (ASL) volume, HCO_3_^−^ concentration and pH caused by dysfunction of the CF transmembrane conductance regulator (CFTR) anion channel ^14–16^. While alteration in ASL volume and physicochemical properties may contribute to abnormal airway mucus in CF, it is worth noting that mucus is also abnormal in internal organs such as pancreas, liver and intestine that have very different luminal environments. An intracellular role for CFTR during mucus maturation and release has previously been dismissed due to the weak or absent CFTR immunofluorescence staining of mucous cells. However, recent studies have demonstrated colocalization of CFTR and mucins in isolated granules from well differentiated bronchial epithelial cells, mRNA transcripts for CFTR and mucins are detected in the same cells using single-cell RNA sequencing, and apparent CFTR immunostaining in ciliated cells may be explained by cross-reactivity of some CFTR antibodies with an antigen in well differentiated ciliated cells ^17–19^. While these findings raise the possibility that CF mucins are abnormal before they are released into the airway lumen, there is currently no direct evidence for a pre-secretory mucin defect.

The viscoelastic properties of complex materials such as mucus are strongly scale-dependent ^4^. Our understanding of mucus/mucin rheology is mostly based on macro- and microscale measurements as mucus has not been investigated at the nanometer (i.e. molecular) scale. Macroscopically, mucus behaves as a viscoelastic gel and its viscosity (defined as the average resistance to flow) is relevant for bulk functions such as mucus transport/clearance and lubrication. However, bulk rheology provides little information on the barrier properties of mucus at the length scales of pathogens and foreign particles ^4^, which is better characterized by micro-rheology techniques such as particle tracking, which measure the viscoelasticity encountered by micrometer-scale entities. Finally, properties at the molecular level need to be addressed using nanometer-scale measurements. For example, nanoviscosity can be used to monitor dissipative forces that oppose molecular motions and are key determinants of protein folding and enzyme catalysis ^20^. Thus, nanoviscosity within compartments of the protein secretory pathway (e.g., ER, Golgi) may influence mucin posttranslational processing and the properties of mucins in storage granules and their rheology after release. Mucin nanoviscosity and its role in disease has not been investigated.

Here, we used a membrane-permeant, BODIPY-based molecular rotor (molecular viscometer) to examine whether CF mucins stored in granules have abnormal physicochemical properties ^21^. We studied nanoviscosity within mucin granules and also in secreted mucus on the surface of well-differentiated primary cultures of CF (F508del/F508del) and Normal (N, i.e., non-CF) human bronchial epithelial (HBE) cells. When combined with fluorescence lifetime imaging microscopy (FLIM), this probe provides reliable estimates of nanoviscosity because the lifetime of the excited state of a fluorophore is independent of its concentration, photobleaching, absorption and excitation intensity ^22^. It reports local (nanometer scale) rotational viscosity, which in intracellular environments is thought to be dominated by water. We validate this assumption, then explore the nanoscale viscosity of mucin molecules before and after release from normal and CF airway epithelial cells.

## Results

### Nanoviscosity within mucin granules

Figure 1A shows representative FLIM images of HBE cells in which mucin granules have been loaded with the BODIPY viscometer. The segmented granules are colored according to their nanoviscosity values measured in centipoise (cP). The noticeable tendency for CF granules to be brighter compared to N granules indicates elevated nanoviscosity. Further, mucin granules in any given cell showed a propensity to have similar nanoviscosity (Figure 1B). A histogram of the median nanoviscosity of granules per cell shows a clear shift towards higher values in CF with a profound right-hand tail representing a larger fraction ∼16% of cells with high viscosity granules (>60 cP) compared to only 2% in N cells, Figure 1C. By contrast, the left side of the histogram of N cells shows a larger subpopulation of cells (∼6 %) with low viscosity (<30 cP) compared to only 1% in CF cells, while the fraction of cells with medium viscosity granules (30-60 cP) are similar for both phenotypes (92% vs 82%), Figure 1D.

**Figure 1.**
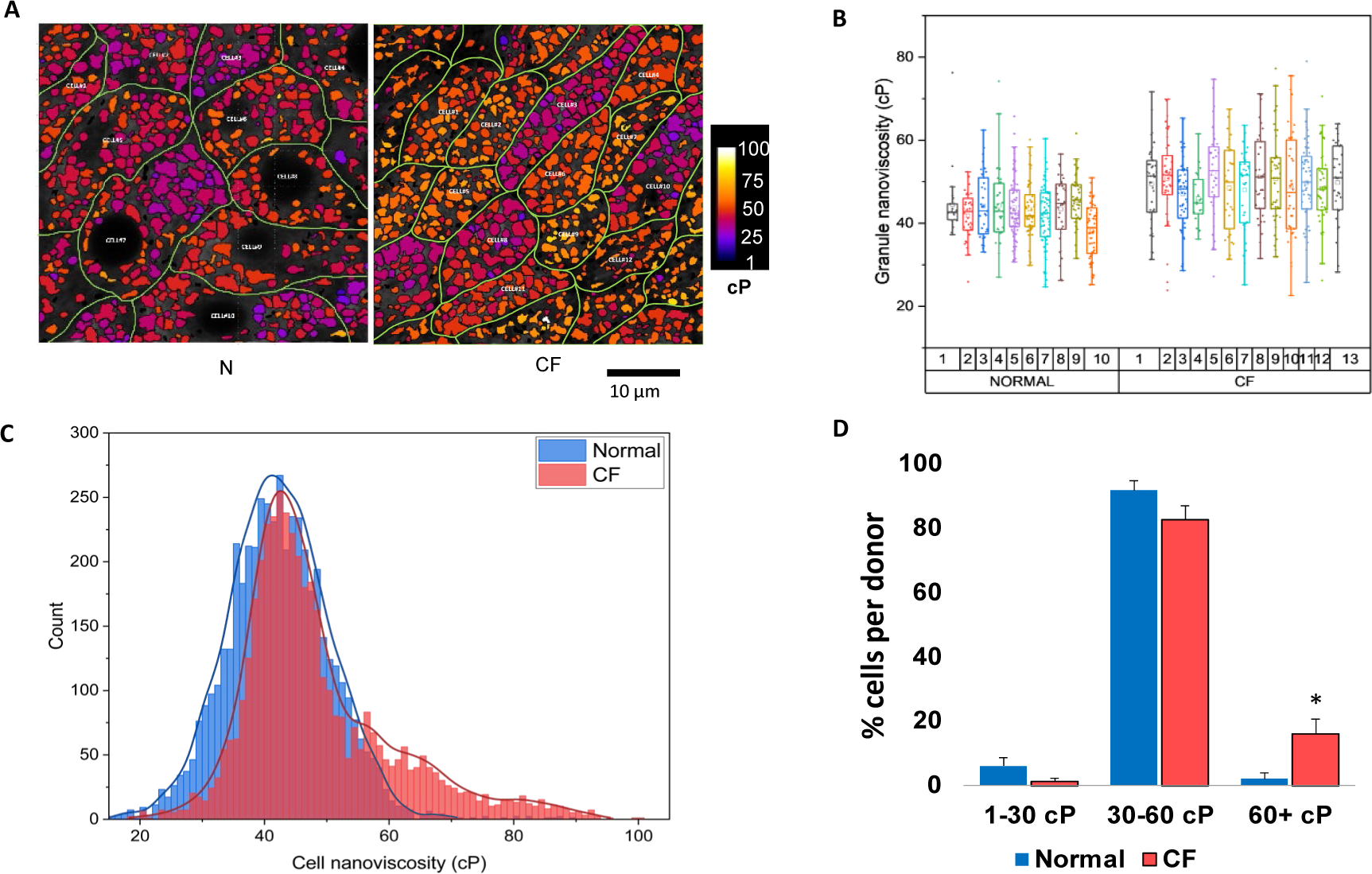
Nanoviscosity of mucin granules measured by FLIM viscometry. **(A)** Representative FLIM images overlaid with color map mask indicating viscosity range of segmented granules in live Normal and CF HBE cells in culture. Nuclei can be clearly recognized as black unstained circles/ovals. Elevated number of orange/yellow-colored granules (indicating elevated viscosity) in CF compared to N cells could be noted. (**B)** Examples of intraluminal nanoviscosity values of granules within individual ten N and thirteen CF cells that were identified as illustrated by their outlines in panel A. Clustering of granules with similar nanoviscosity in separate cells could be noticed. The scatter-box graph also shows a trend of higher median nanoviscosity of granules as well as larger scattering of data in the CF cells compared to N. (**C)** Nanoviscosity distribution of N and CF granules where each count represents median nanoviscosity value per cell. In total 4,888 cells (with 140,102 granules) from 4 N donors and 4,416 cells (with 167,201 granules) from 5 CF donors were analyzed. (**D)** Relative size of cell sub-populations with median granule viscosity per cell per donor falling into arbitrarily assigned ranges: low (1-30 cP), medium (30-60 cP) and high viscosity (60+ cP) in N and CF. Error bars in the graph represent SE, * indicates p<0.05.

Since mucin granule size and its intraluminal viscosity may vary during maturation along the protein secretory pathway, we analyzed the distribution of mucin granule size vs nanoviscosity in N and CF cells. Figure 2 reveals that for both phenotypes, most granules do not show any correlation between their size and viscosity (blue areas in the color-coded density histogram). However, nanoviscosity reaches 200 cP in some CF granules and high viscosity granules (>100 cP) in CF cells tend to be smaller than those in non-CF cells (see the right-hand tail on the histograms).

**Figure 2.**
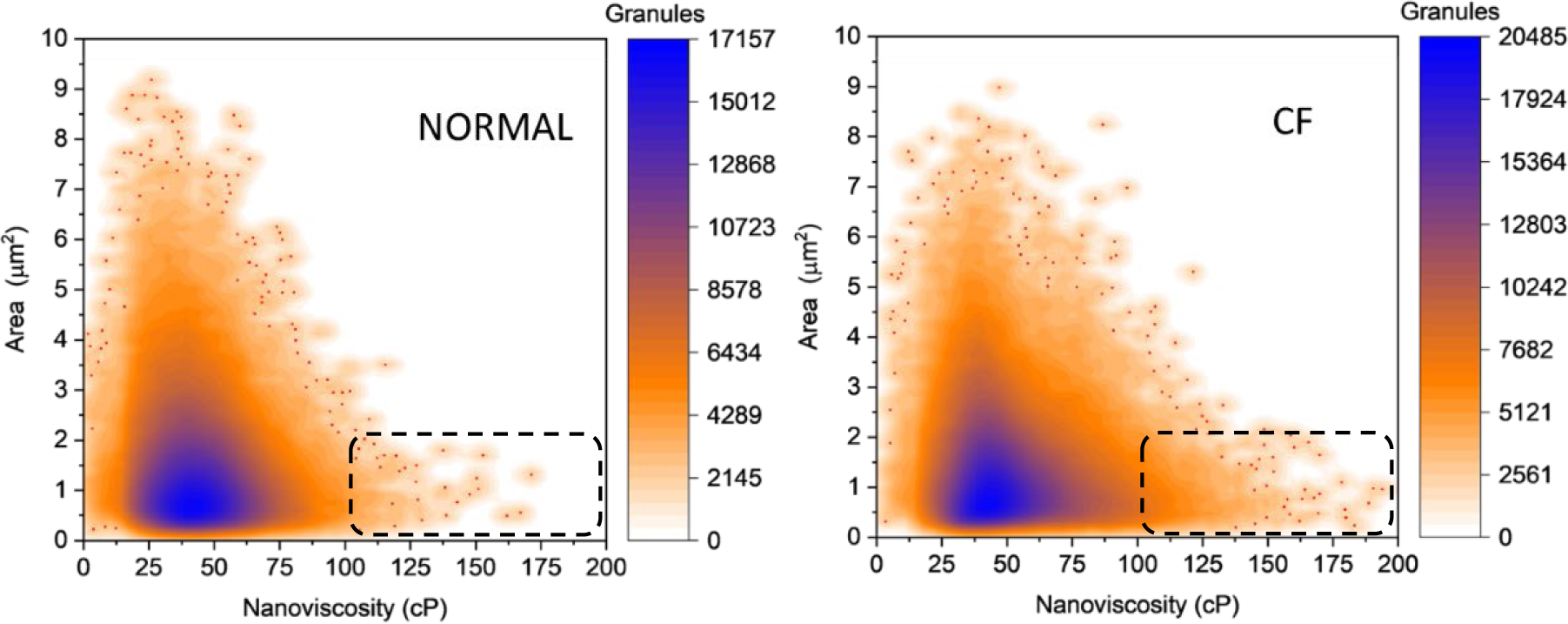
Distribution of mucin granule size (area) vs nanoviscosity in N and CF cells. The color-coded density graph is based on 140,000 granules for N and 167,000 granules for CF cells, respectively. Significantly more small granules with high-viscosity could be seen in CF (indicated by dashed rectangles).

### Secreted mucus nanoviscosity

Abnormally high macro- and micro-scale viscosity of secreted mucus is a characteristic feature of CF ^23, 24^, however viscosity has not been examined previously at the molecular nanometer scale. Here we asked if there is a difference in the nanoviscosity of mucus secreted by N and CF cells when measured *in situ* by adding the diffusible BODIPY-viscometer directly onto the apical surface of HBE cell cultures to avoid potential artifacts due to mucus handling. Figure 3 shows that nanoviscosity, averaged for the entire confocal X-Y plane in the field of view, was 65-70 cP at the cell surface (z=0 µm) for both N and CF cells. By contrast, at z=20 µm above the cells it increased to ∼115 cP on CF cultures while little if any increase was observed above N cells (∼85 cP). Remarkably, the nanoviscosity values for secreted mucus were approximately 2-fold higher than within secretory granules packed with condensed mucins (cellular median of granule nanoviscosity was 40-50 cP). This could not be explained by the altered environment since the probe is insensitive to environmental variables like pH and solvent polarity ^21^. These results suggest that condensed mucins may not be in a highly ordered state within granules (as compared to the expanded and fully hydrated state in secreted mucus), challenging the current paradigm regarding the structural arrangements of stored vs released mucins ^7, 9^. The validity of our mucus nanoviscosity measurements was verified in subsequent experiments.

**Figure 3.**
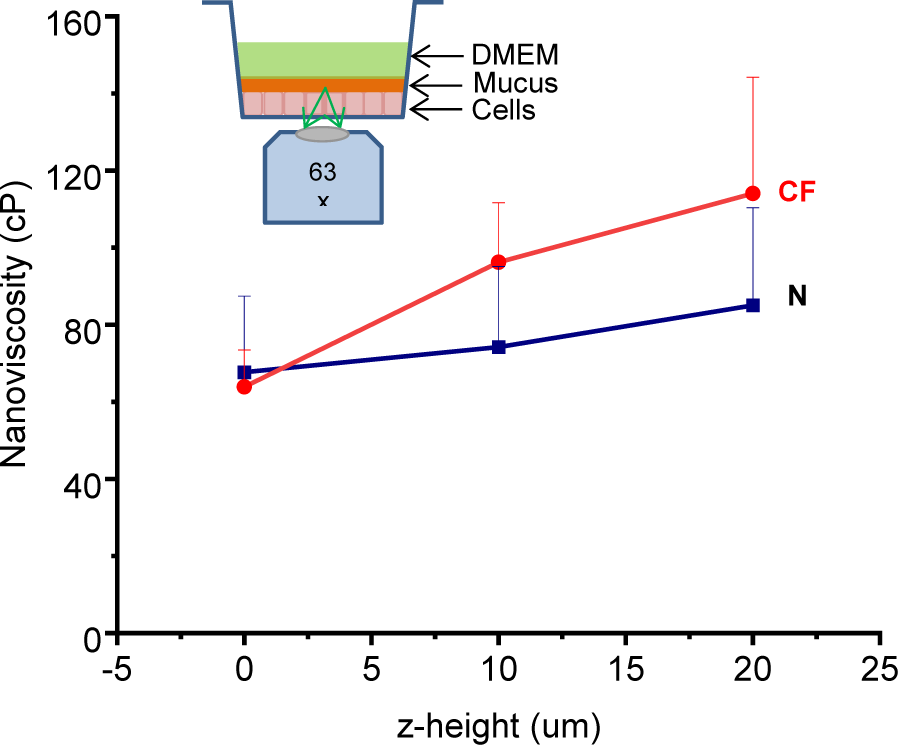
In situ FLIM viscometry of secreted mucus. Each data point represents an average nanoviscosity (+SE) measured for the entire confocal X-Y plane at different z-distance from the top surface of cells. Data is presented for 2 CF donors (6 filters each) and 2 Normal Donors (5 filters each). Cartoon shows cell culture filter and microscope objective of FLIM microscope.

### BODIPY-viscometer reports water nanoviscosity

We asked “What might be the physical basis for the high nanoviscosity of secreted mucus that is reported by the BODIPY-viscometer, and how is it related to the viscosity of water, the major constituent of mucus (up to 98-99%, or ∼55 mol/L) and intracellular environments (60-85%, or ∼40 mol/L)?” The mobility of water is strongly influenced by its interactions with proteins and other macromolecules. When it is constrained in the hydration shell surrounding proteins, water makes an important contribution to the local nanoviscosity that is sensed by proteins and impacts their conformational dynamics and function ^20^. To confirm that the small-molecule BODIPY-viscometer (size ∼1 nm) primarily reports the nanoviscosity of an aqueous phase intracellularly we examined its behavior in well-defined, biologically relevant environments including agarose gels and crude mucin colloid. We found that the BODIPY-viscometer reports the average nanoviscosity corresponding to different water fractions present in the region of interest interrogated by FLIM viscometry. For example, in agarose gels, the fluid phase is dominated by unconstrained water (nanoviscosity ∼1 cP at 20 °C) trapped in large pores of the agarose meshwork (∼100 nm size) (Figure 4A). Thus, nanoviscosity in 2% and 4% agarose gels was indistinguishable from that measured in DMEM medium. By contrast, highly immobilized water (nanoviscosity of 5-45 cP) is detected in the mucin colloid where it is constrained as the hydration shell of mucin macromolecules (Figure 4B). The figure also reveals that the macroscopic viscosity of the same mucin colloid measured in separate experiments using the rolling-ball method correlates tightly with FLIM/BODIPY-measured nanoviscosity, demonstrating that reconstituted mucin viscosity is scale-invariant, as expected for a homogeneous colloid. This agreement also shows that the high nanoviscosity of mucin colloids reported by the BODIPY-viscometer does not result from other interactions, such as binding of the viscometer to mucin chains ^25^. The small size of this viscometer, its ability to accumulate to high concentrations in the mucin granule lumen, and its insensitivity to environmental variables ^21^ makes it well suited for probing the nanoviscosity of intragranular water. These experiments demonstrate that the high intraluminal nanoviscosity in a subset of CF mucin granules results from the presence of highly constrained water, a signature of molecular overcrowding ^20^. It also indicates that the high nanoviscosity of secreted mucus compared to mucins stored within granules is due to the significantly larger fraction of constrained water in mucus compared to granules.

**Figure 4.**
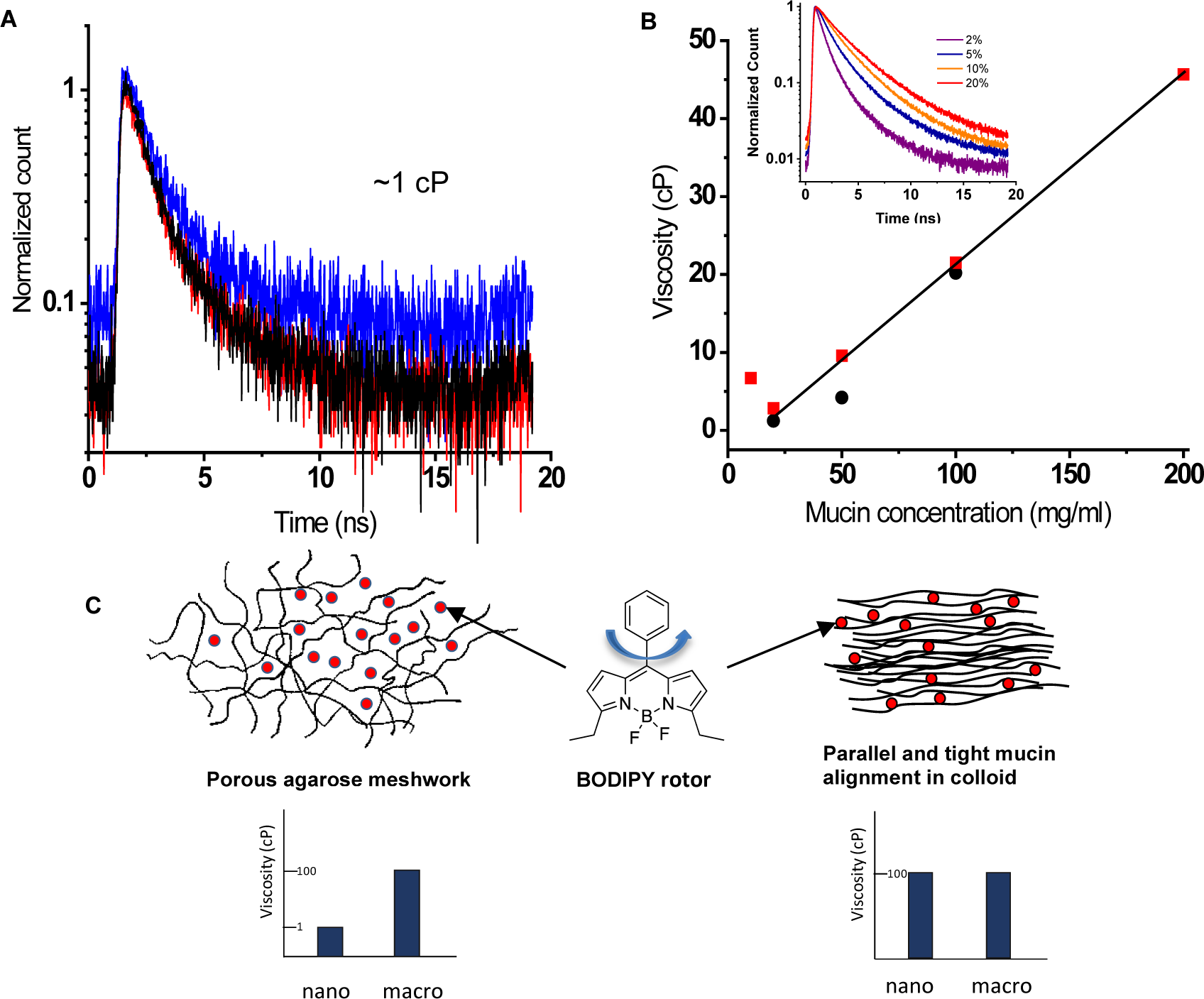
Viscosity measurements of agarose gel and mucin colloid by FLIM viscometry. **(A)** Fluorescence intensity decays of BODIPY viscometer in DMEM cell culture medium (black) and in 2% (blue) and 4% (red) agarose gels. Fluorescence decays were indistinguishable from those observed with DMEM and correspond to nanoviscosity of bulk free water (∼1 cP). This is consistent with a well-structured meshwork of agarose gel where fluid phase is dominated by unconstrained water occupying large pores between agarose polymers, see cartoon below. (**B)** Comparison of mucin colloids nanoviscosity determined by FLIM viscometry (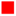) with macro-scale viscosity (●) determined by rolling-ball measurements. Colloids contained different concentrations of porcine stomach mucin Type II reconstituted in 1 N NaOH and pH adjusted to 2 with HCl (see Methods). *Inset:* Examples of fluorescence decay curves for different mucin concentrations (expressed as % of mucins content in the colloid). Crude mucin preparation produced fluorescence decays corresponding to average nanoviscosity higher than that of free water, 3-7 cP for the lowest mucin concentrations of 10-40 mg/ml, and it increased linearly with increasing concentration reaching 45 cP for 200 mg/ml. (**C)** cartoons illustrate proposed structure of agarose gel and mucin colloid, respectively. The corresponding bar graphs illustrates how nanoviscosity compares to macroviscosity for agarose and mucin colloid. The large difference between nano- and macroviscosity in agarose gel, is absent in a homogenous mucin colloid. The latter implies a close contact between mucin polymers in the colloid where, due to steric interactions, they have a tendency to align in parallel.

## Discussion

In this study we found that the average nanoviscosity within the mucin granule (∼20 cP up to 200 cP) is much higher than free water (1 cP), indicating that most intragranular water is constrained by hydrating mucin macromolecules. These values approach those of highly concentrated mucin colloids (40-80 cP, 125-350 mg/ml) and are comparable to nanoviscosities measured previously in the cytoplasm (35 cP), ER lumen (68 cP), and high viscosity intracellular compartments (160-260 cP) using other BODIPY-based viscometers ^25–27^. However, in those reports, high nanoviscosities were not explicitly attributed to the reduced mobility of constrained water. High nanoviscosity is observed in intraorganellar environments involved in protein synthesis and trafficking, where the high concentration of proteins and other macromolecules is expected to cause overcrowding. Macromolecular crowding is also likely in granules, most notably in the large subpopulation (16%) of CF cells that have mucin granules with abnormally high nanoviscosity. This is the first demonstration that CF mucins are abnormal prior to release from the cells. How defective CFTR anion channel alters physicochemical properties of mucins requires further investigation, but it may involve ER stress or reduced salt and water flux into the granules from the cytosol. Single-cell RNA sequencing studies have demonstrated CFTR transcripts in a subpopulation of mucus-secreting cells ^28^ and CFTR protein has been detected by immunofluorescence in the membrane of isolated granules that coimmunostain for mucins^18^.

Altered water homeostasis may lead to molecular crowding in mucin granules. Further, crowding may begin in the ER due to the accumulation of unfolded proteins, leading to ER stress and triggering the unfolded protein response (UPR) to maintain proteostasis. Mucus-secreting cells are particularly sensitive to ER stress due to the enormous size of mucin macromolecules they produce ^29^. Any additional demand for protein synthesis, even in the absence of infection/inflammation may overwhelm the quality control system in vulnerable cells, allowing the misfolded proteins, such as mutated CFTR, to escape ER associated degradation (ERAD) and accumulate in the protein secretory pathway. ER stress can be further amplified by stress-induced overproduction of mucins in CF mucous cells ^30, 31^. Crowding increases water nanoviscosity exponentially due to collective hydration of macromolecules that could extend over 20-40 Å, increasing the nanoviscosity experienced by proteins by an order of magnitude and profoundly slowing their dynamics ^32, 33^. Inhibiting protein movement, folding and enzyme activity is expected to impair the biogenesis of mucin macromolecules and inhibit their glycosylation due to the broad spectrum of enzymes involved in the attachment of oligosaccharide chains. Indeed, increased abundance of weakly charged, under-glycosylated MUC5B has been reported in CF mucus ^34^, although other studies obtained variable results ^30^. However a clear-cut difference may be difficult to establish using conventional methods when abnormally elevated intragranular nanoviscosity is present in a fraction (16%) of CF cells and also occurs in a small fraction (2%) of N cells.

The hydration and lubricating properties of mucins depend on the extent of glycosylation. Removing it reduces water binding by 3.5-fold and increases friction between lubricating mucin layers 100-fold ^35^. Thus, abnormal post-translational processing that leads to altered glycans *in vivo* may yield mucus with reduced lubricating properties ^30^. Earlier observations of impaired mucus exocytosis and swelling in CF cells are compatible with an intrinsic mucin abnormality in CF ^36, 37^. Interestingly, high levels of weakly charged MUC5B glycoforms have also been observed in asthma and in chronic obstructive pulmonary disease (COPD), and in healthy smokers who have an acquired CFTR deficiency, suggesting that CFTR dysfunction may be a common feature of obstructive airway diseases ^1, 34, 38^ with genetic or environmental origins ^39^. Our study suggests that therapeutic approaches that reduce molecular crowding by alleviating ER stress through reduced protein misfolding or protein synthesis and/or enhancing degradation of the unfolded proteins may improve mucus properties ^29, 40^. Reducing the rate of protein synthesis also improves the folding of deltaF508-CFTR ^41, 42^.

Secreted mucus had a nanoviscosity of 65-115 cP when measured *in situ*, up to twice the median of intra-granule values (40-50 cP). This indicates nearly all water surrounding the probe in extracellular mucus is highly constrained, in contrast to agarose gels, where large amounts of free water are within pores in the matrix. The close association between mucin polymer and water differs from the classical view of airway mucus as a porous agarose-like hydrogel ^9^. However, if contact between extracellular linear mucin chains, such as that of MUC5B ^43^ occurs in a parallel ordered arrangement, it may lead to formation of nematic liquid crystalline sheets ^44^ (Figure 4C), analogous to the crystalline protein structure of the eye lens ^45^. Liquid crystalline microstructures have been observed in mucus from human trachea and from a slug using polarization microscopy ^7, 44^. In the airways they may produce a laminated morphology that allows low-friction axial sliding/flow over the epithelial surface propelled by beating cilia while providing a physical barrier to transverse penetration by pathogens, Figure 5. Our finding that mucus nanoviscosity increases with distance from the epithelial surface supports this view of stratified mucus layers that have increasing nanoviscosity. Elevated mucus nanoviscosity in CF (115 cP) compared to N cultures (85 cP) may reflect under-glycosylation that results in a smaller quantity of bound water (i.e., reduced water-binding capacity), a smaller volume, and increased protein concentration as observed previously, e.g., in ref.^46^. It also increases mucin friction, favors aggregation/entanglement, and may lead to formation of more mucin strands and impaired mucociliary transport as reported previously ^47^. Although we observed slightly higher mucus viscosity in CF compared to N, it remains far from the differences reported in the literature, which reach > 1000-fold ^23^. The initial viscosity difference, while small at the beginning, would likely trigger a series of diverging events including, mucus accumulation, mucostasis and infections, leading to much larger viscosity differences over time between CF and Normal due to, e.g., increased impurity from dead cells and pathogens.

**Figure 5.**
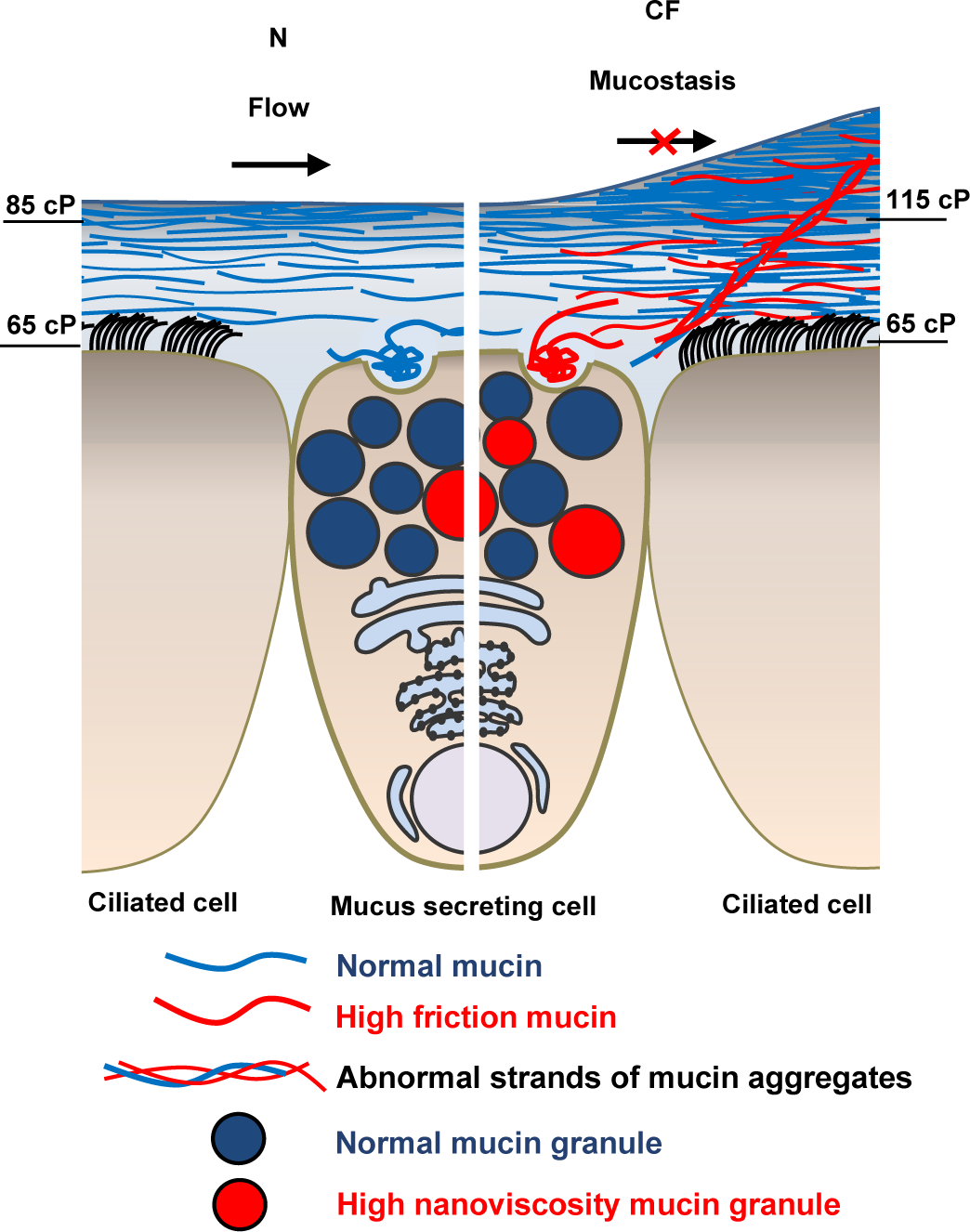
Stratified arrangement of mucus layer. Simplified diagram of airway surface epithelium with mucus layer (not showing submucosal glands). Aligned in parallel mucins such as linear MUC5B form a liquid crystalline structures permitting low friction axial sliding propelled by beating cilia. In CF, fraction of underglycosylated high-friction mucins, and their aggregates forming mucin strands, perturb mucociliary transport, leading to mucus accumulation and mucostasis. Mucus nanoviscosity is much higher compared to free water and increases towards top of mucus layer from 65 cP (for N and CF, z=0) to 85 cP in N and 115 cP in CF (z=20 µm).

An increase in nanoviscosity after mucin secretion implies that intragranular mucins are not uniformly structured despite their packing density. This finding is difficult to reconcile with the block copolymer model ^7^ and concatenated rings model for intestinal mucin MUC2 ^9^, which posit tight and ordered packing, Figure 6. Several factors could lead to compaction without highly ordered structure. Unlike secreted mucus, intragranular mucins are not fully hydrated and occupy less volume, and macromolecular crowding itself promotes compaction, conformational collapse and aggregation ^20^. Intragranular mucins may also condense spontaneously into random coils that form globular particles as condensation of mucins into nano-size particles can be induced *in vitro* by glycerol and is reversible ^48^, suggesting there is minimal knot formation and entanglement of the mucin chains contrary to previous assumptions ^7, 9^. Rather, polyanionic mucins may behave like “slippery spaghetti” due to repulsive electrostatic forces between the mucin’s polymer chains. Finally, elevated intragranular Ca^2+^ and H^+^ concentrations promote the condensation of intragranular mucins ^6, 49^, which occurs when mucins are in disordered random coils independently of their topology. Our data are most consistent with a scheme in which intragranular mucins form a weakly-ordered colloid of mucin macromolecules as collapsed random coils and their aggregates, possibly with mobile and immobile fractions^11, 50, 51^.

**Figure 6.**
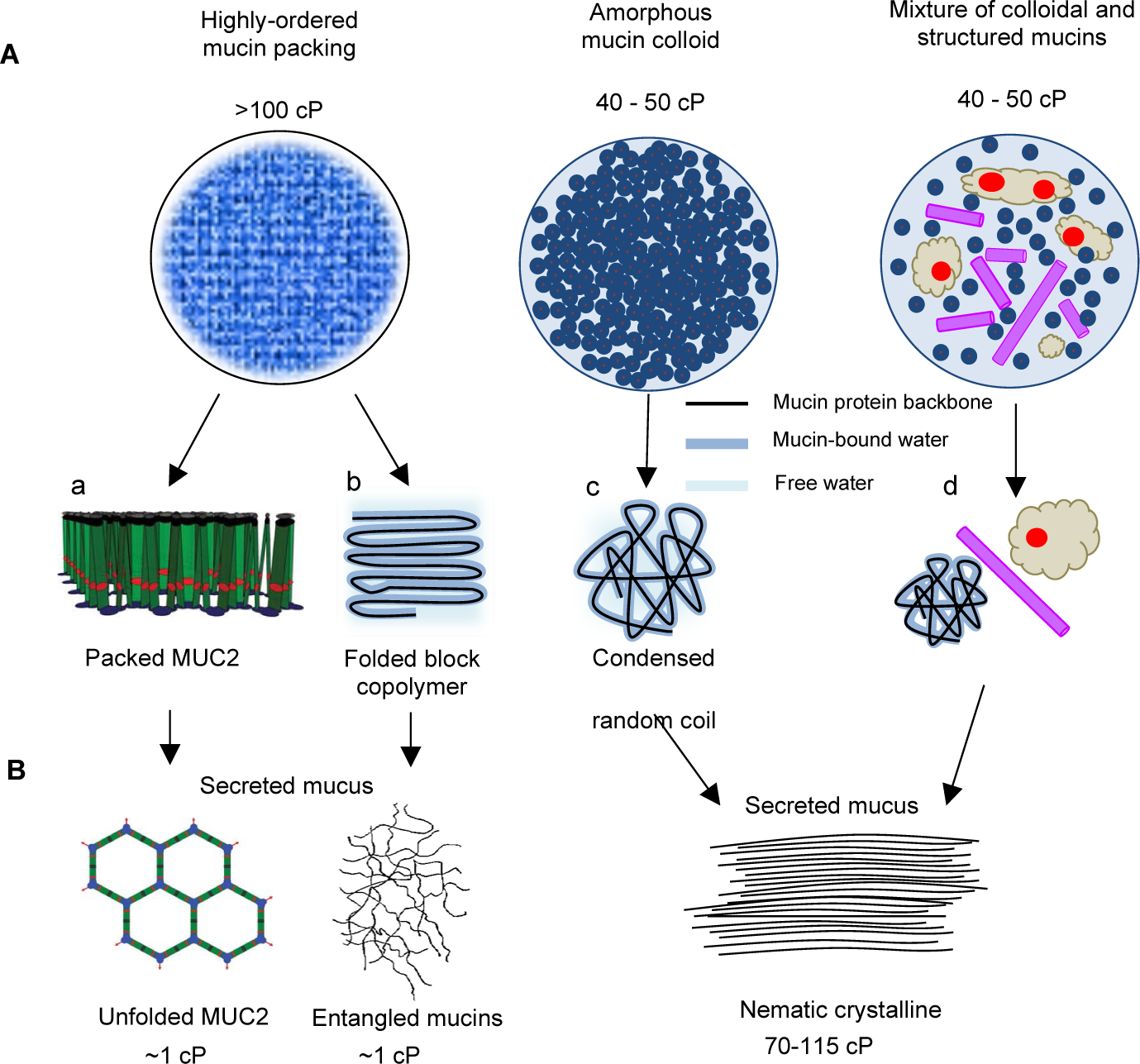
Proposed structure of intragranular mucins and secreted mucus. **(A)** Current models for intraluminal mucin granule organization include: a) well-organized MUC2 packaging ^9^, and b) crystalline array of folded block copolymer ^7^. Both models imply well-ordered intragranular mucin structure consistent with most water in a highly constrained state, expected nanoviscosity ≥100 cP and little free water. This study, model c), suggests weakly-ordered colloidal state of compacted random coil mucins, or d), a mixture of random coiled and distinctly structured mucins, such as disk/filament-like and electron-dense matrix ^52^, consistent with significant fractions of free and bound water leading to moderate intraluminal nanoviscosity (median 40-50 cP). (**B)** Secreted mucus structure proposed by different models. MUC2 unfolded net-like network and entangled mucin matrix, model a) and b), imply large spaces between mucin polymers filled by free water (∼1 cP nanoviscosity), similar to agarose gel, see Figure 4C. This study finds high mucus nanoviscosity (70-115 cP) suggesting nematic liquid crystalline structures where majority of water is highly constrained in the hydration shell of aligned in parallel linearized mucins.

Moderate intraluminal granule nanoviscosity is also expected when mixtures of colloidal mucins and their condensates undergo organized intragranular folding/restructuring and spatial segregation during granule maturation ^52^. Intragranular nanoviscosity appears to be a well-controlled parameter despite increasing granule size during maturation. It remains moderate (25-75 cP, blue areas on the density graphs, Figure 2) for vast majority of granules and does not show any correlation with size. The large fraction of small granules with high nanoviscosity (>100 cP) suggests there was less maturation of mucins in CF cells.

### Conclusions

The significant elevation of intraluminal nanoviscosity in a population of CF mucin granules found in this study demonstrates there is an inherent, pre-secretory mucus defect in CF. The high nanoviscosity of secreted mucus compared to mucin granules implies close contact between the polymers that could be only achieved by nematic crystalline-like structures in which most mucins have a parallel alignment. Such alignment would lead to a stratified mucus structure rather than a porous agarose-like gel. Moderate nanoviscosity within mucin granules (approximately 2-fold lower than in the secreted mucus), implies a weakly-ordered structure rather than highly-ordered tight packing. It suggests that most stored intragranular mucin chains form a weakly-ordered colloid as collapsed random coils and their aggregates.

High nanoviscosity of intracellular CF mucins suggests molecular overcrowding that is expected to impair their post-translational processing, resulting in abnormal mucus glycosylation, reduced water-binding capacity, and diminished lubricating properties. This mechanism is likely to be an important factor in CF mucus pathology and may provide a unifying explanation for abnormal mucus in different CF-affected organs, and in COPD where there is an acquired deficiency in CFTR due to cigarette smoke exposure. Our study suggests new therapeutics that target early steps in the mucin secretory pathway may be useful for the treatment of muco-obstructive lung diseases.

## Methods

### Donors

CF lung tissues from five F508del/F508del patients (median age 30 years) were obtained from the Montreal Clinical Research Institute (IRCM) with informed consent after lung transplantation following protocols approved by the Institutional Review Boards of the IRCM and McGill University (#A08-M70-14B). N (non-CF) lung tissues were obtained from four donor organs not employed for transplantation (median life span 59.5 years).

### Human bronchial epithelial cells

Normal and CF primary HBE cells were prepared in the Primary Airway Cell Biobank (PACB) of the CF Translational Research Center (CFTRC, McGill University, Montreal, QC), as previously described ^53^. Briefly, freshly isolated cells were seeded on flasks coated with PureCol (Cedarlane Laboratory, Burlington, ON, CA) and cultured in CnT-17 medium (CellnTec Advanced Cell Systems, Bern, CH) until 80% confluence was reached. Normal or CF HBE cells were then detached with trypsin solution, seeded on permeable 12 mm Transwell® 0.4 µm Pore Polyester Membrane Inserts (Corning, NY, USA) coated with collagen IV (Sigma-Aldrich, ON, Canada) and cultured in CnT-17 until confluency. Next, the apical medium was removed to create an air–liquid interface and the basolateral medium was replaced with differentiation medium (1:1 volume of BEGM and DMEM (Life Technologies, CA, USA) supplemented with 1.5 µg/mL BSA, 0.1 µM retinoic acid, and 100 U/mL of penicillin-streptomycin every two days for at least 35 days to obtain highly differentiated cultures ^54^.

### Histology

Hematoxylin and Eosin (H&E) and Alcian Blue-Periodic Acid Schiff (AB-PAS) staining of HBE cells were performed by Histology Core Facility at McGill Goodman Cancer Research Centre following standard methodology. Images of histology slides were acquired with an Olympus BX53 microscope (Olympus, Japan) and an Exi Aqua bio-imaging camera (Exi Aqua, CA). Histology was performed for each batch of cultured HBE cells used in the study. Typical histology micrographs of N and CF HBE cells prepared by the facility could be found in ^18^.

### Fluorescence lifetime imaging microscopy (FLIM) of HBE cells

We used a previously described BODIPY molecular viscometer to visualize and quantify the viscosity of mucin granules in live HBE cells after differentiation at the air-liquid interface ^21^. For imaging, cell inserts were cut from their plastic supports and incubated in Gibco’s phenol red-free DMEM (Invitrogen Canada, Burlington, ON, Canada) solution containing bicarbonate and 2.3 µM BODIPY-viscometer for 40 min at 37°C, 5% CO_2_. The inserts were washed with DMEM and placed in an experimental imaging chamber (Warner Instruments, LLC, Hamden, CT) containing ∼150 µl DMEM solution. In these experiments the apical side of cells faced the bottom coverslip (25-mm diameter glass #1). The chamber was mounted on the stage of a Zeiss AxioObserver (Zeiss, Jena, Germany) motorized inverted microscope with control of temperature (37°C) and CO_2_ (5%). Fluorescence lifetime images of HBE cells with visible mucin granules were acquired using a Zeiss LSM 710 confocal imaging system and a 63×1.4 NA plan apochromat oil immersion objective (Zeiss). The microscope was outfitted with a PicoHarp 300 controller (PicoQuant, Berlin, Germany) consisting of a FLIM excitation source, an internal laser bypass and a single-photon avalanche diode detector. A 473 nm pulsed laser (PicoQuant LDH series) operating at 50 MHz pulse rate served as an excitation source. Fluorescence emission was collected after passage through an Alexa Fluor 488 nm filter. The pinhole aperture was set at 43 µm and scans were acquired during ∼150 s in fast scan mode (0.8 s per scan). The size of acquired scans was 512×512 pixels for a physical area of 45×45 µm. Scans were recorded with SymPhoTime software (PicoQuant) and stored as Pt3 files.

For *in situ* mucus viscosity measurements, 400 μl of DMEM solution containing 11.5 µM BODIPY-viscometer was added on apical side of HBE cells grown on permeable filters and mounted immediately on the stage of the microscope (see Fig. 3). Experiments were performed at ∼30°C. Scans were acquired at several planes starting from near the top of the epithelium (apical surface) and moving towards the surface of the mucus layer in displacement increments of 10-15 µm. When applicable, scans from different planes of acquisition were combined in order to increase the signal to > 1,500 counts. Data from 2 CF cells donors (total of 12 filters) and 2 N cells donors (total of 10 filters) were analyzed.

### Image processing

The FLIM scans stored in Pt3 files were converted into stack of 512×512 TIFF images (8-bit) using a modified version of the publicly available ImageJ Java plugin Pt3Reader (June 7th, 2016 version; https://imagejdocu.tudor.lu/doku.php?id=plugin:inputoutput:picoquant_.pt3_image_reade r:start). The plugin was modified for compatibility with our Pt3 files and to limit the size of stack files by keeping only the initial, stronger part of the signal (0-19 ns). The time resolution per slice was 16 ps. Each pixel value in each slice represented the sum of the photon event count in all scans for this pixel location and corresponding time. The intensity image was generated by summing all time slices into a 32-bit TIFF image. This intensity image was then segmented with the ImageJ plugin Weka trainable segmentation ^55^. To further increase detection of the granule contours we applied the built-in ImageJ Watershed segmentation algorithm. Finally, granule regions of interest (ROIs) were created from the segmented images. Only ROIs having >25 pixels and meeting a circularity criterion within 0.25-1.0 were kept for analysis.

### Lifetime determination and nanoviscosity colormap

In our previous report ^21^, the fluorescence lifetimes (LT) of individual granules were extracted from the Symphony^TM^ software-generated Fast_LT images, in which each pixel is assigned a LT level by subtracting the barycentre of the instrument response function (IRF) from the barycentre of the pixel decay signal. Instead of using this rapid approximation approach, we implemented in this study a more robust LT determination described below. Combining this improved method with revised calibration data improved the accuracy of granule viscosity determinations compared to our previous study ^21^.

For LT determinations, the photon count over time (1,200 slices, 0-19 ns) was compiled for each ROI (1 ROI per granule) and the exponential fluorescence decays were calculated for each granule from the photon count trace over a fixed time interval (0.7 to 6.1 ns past the peak). The trace collection and LT calculation were automated using a custom-made ImageJ macro. The LT was computed with a simple linear regression fit to the natural logarithm of the photon count values. Only LT with R^2^>0.6 were kept for downstream analysis and then converted into nanoviscosity. For conversion of LT to nanoviscosity, two distinct calibration functions for separate LT ranges were used. For LT≤2.7 ns (or η ≤ 15 cP): η (cP) = 0.85 + 2.65×10^−5^×e^(4.85×LT)^ where LT is in nanoseconds and for LT>2.7 ns (or η > 15 cP): η = 10^(0.70×LT-0.70)^. Most granules in this study had LT>2.7 ns.

### BODIPY-viscometer calibration

Calibration experiments of the BODIPY-viscometer molecule were performed in biologically relevant environments such as agarose gel and crude mucin colloid.

Agarose gels of 2 and 4% w/vol of DMEM were prepared according to standard procedures. After solidification (>24 hours), a small block of gel ∼10×5×2 mm was cut and a drop of ∼50 µl of DMEM solution containing 2.3 μM BODIPY probe was deposited on the surface of the block. Gel blocks were incubated for at least 15 minutes to allow the probe solution to diffuse throughout the gel, then placed inside the chamber on the stage of the FLIM imaging system (∼30°C).

BODIPY probe fluorescence intensity decay curves were also recorded in colloid solutions containing different concentrations (10, 20, 50, 100 and 200 mg/ml) of porcine stomach mucin Type II (Sigma-Aldrich, ON, Canada). Mucins isolated from porcine stomach were dissolved in 1 M NaOH and the pH subsequently adjusted to ∼2 with HCl ^56^.

### Rolling ball viscosity measurements

To validate FLIM measurements of mucin colloid viscosity we performed parallel macroscopic viscosity measurements using a capillary and small ball drop device as described previously ^57^. We assessed the macroviscosity of colloid solutions containing 20, 50 and 100 mg/ml porcine stomach mucin Type II prepared as described above (*BODIPY-viscometer calibration*). For higher mucin concentrations, exceeding 100 mg/ml, macroviscosity of colloid is too high and could not be determined by rolling ball method.

### Data organization for analysis

We categorized the granule data into two nested levels: Phenotype and Cell. The two Phenotype groups (N and CF) pooled all individual granules irrespective of other identifiers. The Cell ensembles contained the information for all granules at their particular level of integration while being also divided according to their phenotype. A custom ImageJ macro generated the normalized distribution for all categorized sets and calculated their median.

## Author Contributions

O.P. and F.B performed imaging experiments and data analysis. S.V.D. synthesized BODIPY-viscometer. I.G., Z.G., R.F. and S.V.D. advised on dye application for cell imaging. F.B., O.P. and R.G. designed experiments with input from I.G., Z.G., R.F. and J.W.H. F.B., O.P. and R.G. prepared figures. F.B., R.G. and J.W.H. wrote the manuscript with input from O.P., Z.G. and S.V.D.

## Notes

The authors declare no competing financial interests.

## Acknowledgements

The authors thank the Primary Airway Cell Bank (PACB) of Cystic Fibrosis Translational Research Center (McGill University) for providing HBE cell cultures. CF lung tissue was obtained with CF Canada financial support from the Respiratory Tissue and Cell Biobank of the CRCHUM (Centre de recherche du Centre Hospitalier de l’Université de Montréal), which is directed by Dr. Emmanuelle Brochiero and affiliated with the Québec Respiratory Health Research Network. FLIM images were collected in the McGill University Advanced BioImaging Facility (ABIF), RRID:SCR_017697”. We thank Tsung Huang (Faith) for performing rolling ball viscosity measurements with reconstituted mucins (Fig. 4B).

